# Lineage space and the propensity of bacterial cells to undergo growth transitions

**DOI:** 10.1101/256123

**Authors:** Arnab Bandyopadhyay, Huijing Wang, J. Christian J. Ray

## Abstract

The molecular makeup of the offspring of a dividing cell gradually becomes phenotypically decorrelated from the parent cell by noise and regulatory mechanisms that amplify phenotypic heterogeneity. Such regulatory mechanisms form networks that contain thresholds between phenotypes. Populations of cells can be poised near the threshold so that a subset of the population probabilistically undergoes the phenotypic transition. We sought to characterize the diversity of bacterial populations around a growth-modulating threshold via analysis of the effect of non-genetic inheritance, similar to conditions that create antibiotic-tolerant persister cells and other examples of bet hedging. Using simulations and experimental lineage data in *Escherichia coli*, we present evidence that regulation of growth amplifies the dependence of growth arrest on cellular lineage, causing clusters of related cells undergo growth arrest in certain conditions. Our simulations predict that lineage correlations and the sensitivity of growth to changes in toxin levels coincide in a critical regime. Below the critical regime, the sizes of related growth arrested clusters are distributed exponentially, while in the critical regime clusters sizes are more likely to become large. Furthermore, phenotypic diversity can be nearly as high as possible near the critical regime, but for most parameter values it falls far below the theoretical limit. We conclude that lineage information is indispensable for understanding regulation of cellular growth.

**Author Summary:** One of the most important characteristics of a cell is whether it is growing. Actively growing cells can multiply exponentially. In the case of infections and cancer, growth causes problems for the host organism. On the other hand, cells that have stopped growing can allocate cellular resources toward different activities, such as bacteria surviving antibiotics and tissues in multicellular organisms performing their physiological roles. Observing small bacterial colonies in a microscope over time, we have found that cells closely related to each other often have similar growth state. We were curious if lineage dependence was an intrinsic property of growth regulation or if other factors were needed to explain this effect. We therefore built a computational model of a growing and dividing cellular colony with an encoded growth regulation network. We found that regulation of growth is sufficient for lineage dependence to emerge. We next asked if lineage dependence constrains how diverse the cellular population can become. We found that cellular diversity can reach a peak that is nearly as high as possible near the conditions that have the highest lineage dependence, but that most conditions do not permit such high diversity. We conclude that lineage is an important constraint and discuss how the growth arrest transition is in some ways like a phase transition from physics, and in some ways strikingly different, making it a unique phenomenon.

## Introduction

The process of cellular growth is both the distinguishing feature of living matter and central to the roles of regulatory networks from microbes to metazoa. Growth and division is also a primary source of phenotypic diversification. For instance, when a bacterial cell divides, and its cellular contents become partitioned into two daughter cells, diffusible cytoplasmic components are often randomly distributed into the daughter cells in a binomial distribution. Such phenotypic diversification permits populations to be robust to unpredictably changing environments, a phenomenon known as bet-hedging. A striking example of this effect is the regulation of growth rate by toxins.

Most of the molecular content in the bacterial cytoplasm undergoes growth-mediated dilution (in some cases, such as most proteins, as the primary mechanism of degradation). Reduction in cellular growth rate by a cytoplasmic toxin, or other molecule with toxic effect, creates an effective positive feedback loop, trapping some cells in a growth arrested state until they can escape in changed conditions [1-3]. This mechanism is associated with antibiotic-tolerant persister cells arising in the population, which cause difficulty in antibiotic treatment [4]. Various feedback mechanisms are associated with growth bistability [5]. Thus, understanding the processes that result in growth diversification is an important goal on the path to solving the impending antibiotic resistance crisis.

Growth arrested cells typically represent a small subset of a bacterial population [6]. In *E. coli*, growth arrested persister cells are associated with alterations in metabolic activity via the stringent response [7, 8], and with efflux of antibiotics [9]. Depending on the mechanism of induction, persister cell fractions can be spontaneously produced or respond to external stresses [6]. Persistence in *E. coli* is associated with toxin-antitoxin systems and global metabolic regulation [10], with a core mechanism of toxins that are neutralized by antitoxins [11] (Fig. 1a-b). The competing effects of toxin and antitoxin create a threshold in a stoichiometric effect via molecular titration that can cause conditional cooperativity of TA gene regulation [12, 13]. When accounting for gene expression noise and proteolysis of antitoxins, free toxin levels will gain sufficient concentration to result in a growth feedback mechanism that ultimately induces growth arrest in above-threshold cells. The result is skewed phenotypic distributions, with a core fast-growing group of cells along with rarer, growth arrested cells, as opposed to regression to mean levels observed in networks without the growth arrest threshold (Fig. 1c-d).

**Figure 1.**
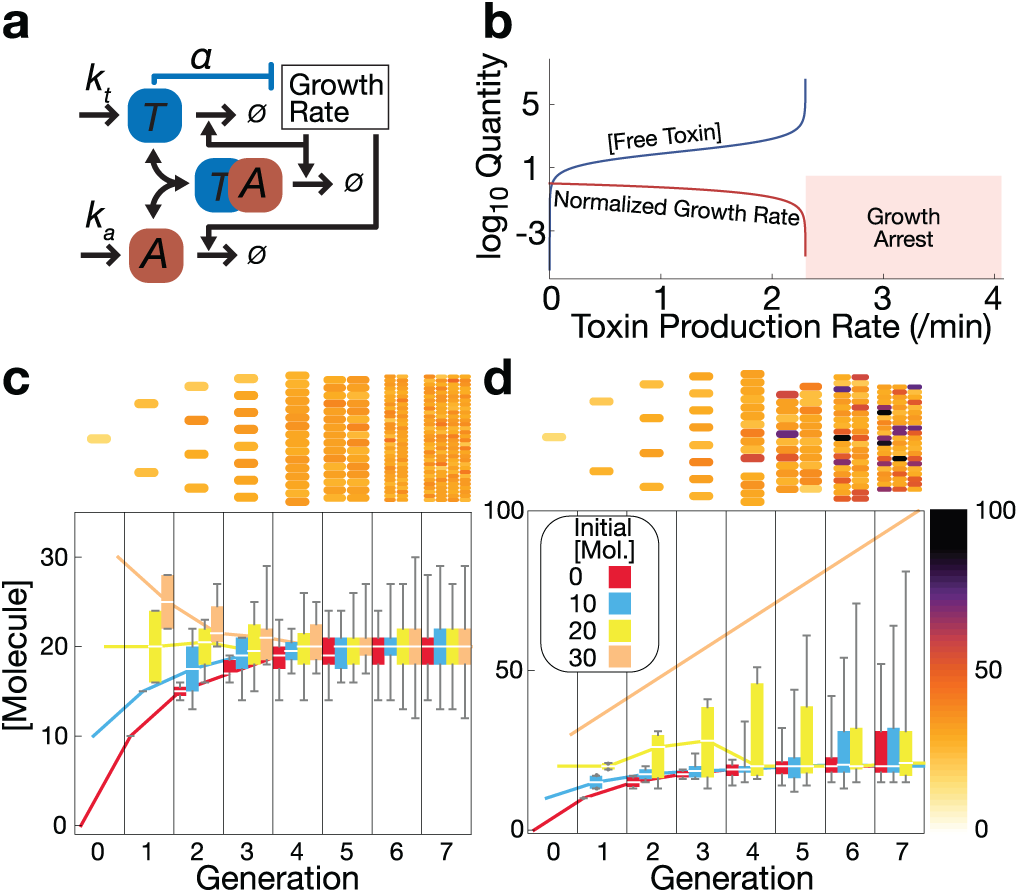
Simulated effects of a molecular network with an endogenous growth-regulating threshold in bacteria. **a**. Simplified toxin-antitoxin module, depicting its interaction with cellular growth rate. **b**. Deterministic steady state model predictions for a toxin with growth feedback. A regime with no deterministic molecular steady state (labeled “Growth Arrest”) arises when toxin production sufficiently exceeds the growth feedback-imposed threshold. Growth rate is normalized to the maximum = 1. **c**. Binomial phenotypic inheritance at a constant molecule production rate. With no effect on cellular growth rate, the population exhibits regression to the mean within a few generations of division. **d**. With a discrete growth arrest threshold, the population becomes increasingly skewed over time. Box and whisker plots represent median, interquartile range, and range of a population started from a single simulated cell. Details on model implementation are presented in Supplemental Materials.

Motivated by observations on phenotypic inheritance [14-16] and the effects of lineage correlations on daughter cell phenotypes [17-21], we asked how much phenotypic diversity could be attained for various levels of endogenous growth regulation, and to what extent lineage determines phenotypic outcomes. Based on our previous study [17], we hypothesized that a higher chance of growth arrest amplifies the effects of cellular lineage on phenotypic correlations.

To explore this hypothesis, we used an established experimental model of threshold-based growth arrest in *E. coli* to experimentally confirm lineage dependence. We then created a minimal multiscale computational framework that allowed more extensive characterization of the various growth regimes than were possible with time-lapse microscopy. Our computational model represents the processes of cellular growth and division, with binomially distributed inheritance of a simplified toxin-antitoxin-like system subject to stochastic molecular kinetics in individual cells over time. We modeled a functional dependence of growth on toxin concentrations as an exponential function with a key parameter, *α*, that quantifies how toxic the toxin is. We used various specific realizations of the framework to simulate growth of small bacterial populations from a single common ancestor and growth regulation by the simulated toxin for various toxin:antitoxin production ratios. Our computational results confirm and extend the experimental results, showing that the bet-hedging regime results in complex lineage structures.

These results show, for the first time, how important lineage is to growth regulation and bet-hedging phenotypes involving growth. Consideration of lineage is now indispensable for studies on phenotypic heterogeneity, phenotypic memory, and regulation of the growth arrest transition. Finally, our results suggest that lineage space used in evolutionary [22] and multicellular organism development studies [23] is an important concept to apply in studies of bacterial phenotype.

## Results

### Lineage Dependence in an Experimental Model

We first sought to establish an empirical basis for growth arrest kinetics and threshold-based amplification of lineage correlations. An established experimental model of threshold-based growth arrest [17] provided a simple way to track growth in a lactose-sensitive strain of *E. coli*. In this model, lactose stimulates growth at sufficiently low concentrations, but creates toxicity in a subset of cells at high concentration that results growth arrest or death of those cells. Presently, the precise mechanism of toxicity is not known in this model, but the competing effects of lactose import rate and processing rate are the most likely culprit, and the threshold-based mechanism for growth arrest and persistence is established [17]. In the high-lactose condition, bacterial colonies have a slow net growth rate and a high likelihood of any individual cell eventually undergoing growth arrest and/or death.

We used time-lapse fluorescence microscopy to track individual microcolonies in a microfluidic device with constant perfusion of fresh minimal medium containing defined concentrations of a single sugar as the sole carbon source. We used two carbon sources: a growth-arrest-prone condition with a high lactose concentration (50 g/l), and a condition that does not induce a growth arrest threshold, with a moderate glucose concentration (2 g/l) (Fig. 2; Movies S1 and S2). As inferred from extension of cellular major axis length, cells grow exponentially at heterogeneous rates (Figs. 2a-b, 2e-f, S1) and are capable of quickly shifting between growth rates, e.g., from fast to slower or non-growing (Fig. 2b, 2f). To identify cases of mid-cell cycle shifts in growth rate, we fit each cell cycle to an exponential growth model, applied Bonferroni correction to the resulting fit significance levels, and selected the non-significant cases (Fig. S3). A constitutive fluorescent reporter provides clear visual evidence of mother-daughter cell correlations only in the growth arrest-prone condition (Fig. 2c, 2g).

**Figure 2.**
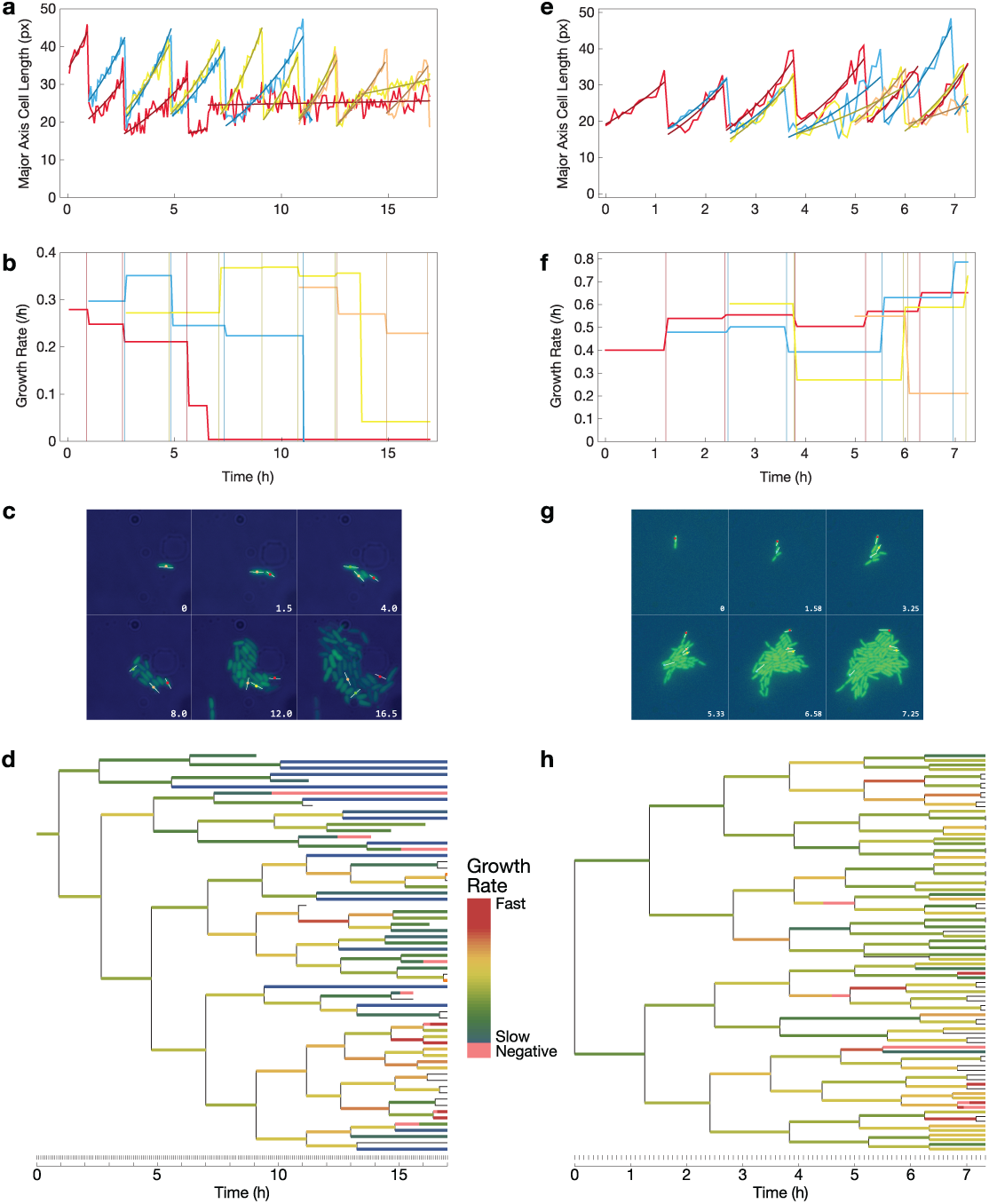
Growth rate of *E. coli* B REL606 GFP+ cells prone to stochastic growth arrest in high lactose reveals lineage dependence. Numbers indicate time in hours. **a** – **d**. Col-ony grown in a commercial microfluidic device with continuous perfusion of minimal medium containing 50 mg/ml lactose as described in Methods. **e** – **h**. Colony grown with continuous perfusion of minimal medium containing 2.5 mg/ml glucose, which does not predispose cells to growth arrest. **a, e**. Growth kinetics of a selection of cells. Individual trajectories are divided by cell division or different growth rates by a leastsquares fit of the data to the model L(*t*) = *L*_0_*e*^*gt*^. **b, f**. Growth rates from exponential model fit. Vertical lines indicate cell division times for the corresponding trajectory color. **c, g**. Selected frames of the time-lapse microscopy experiment. **d, h**. Lineages derived from time-lapse microscopy. Colors indicate growth rate. Lack of color indicates insufficient data for a significant fit. Note asymmetry in **d** and symmetry in **h**.

We reconstructed the microcolony lineage in both conditions to quantify the effects of non-genetic inheritance in this experiment (Fig 2d, 2h). The result of the growth arrest threshold is a striking effect on the structure of the lineages. The growth arrest-prone lineage shows distinct clusters of growth arrested or dead cells, and clusters of faster growing cells, resulting in an asymmetric tree (Fig. 2d). On the other hand, absent the growth arrest threshold, the tree is nearly symmetric (Fig. 2h). In the growth arrest prone condition, we classified cells into being growth arrested or dead (apparent growth rate = 0) or actively growing. Of the 63 total cells in the final lineage, 16 (25.4%) were determined to be completely growth arrested or dead at the final time point. We determined the pairwise lineage distance, defined as the time since the most recent common ancestor, for three subsets: all cells, only growing cells, and only growth arrested cells (Fig. S2). The all-growing and all-growth arrested subsets both had significantly closer lineage distances compared to the all cells set (*p* < 0.05, Mann-Whitney U). From these results, we conclude that lineage has a strong effect on phenotypic heterogeneity during colony development around a growthmodulating threshold.

### Lineage Dependence is Reproduced in a Simple Computational Branching Process Model

To determine the minimal set of mechanisms necessary to reproduce the interactions between threshold-based molecular regulation of growth rate and population dynamics, we created a computational model containing cell agents growing and dividing at a typical rate for enteric bacteria (30 minute doubling time), each with a cell volume and division upon doubling of the volume. Each cell agent has embedded stochastic kinetics of a growth-inhibiting molecule (analogous to a toxin) and a neutralizing molecule that binds and prevents toxicity (analogous to an antitoxin). As discussed in more detail in Methods, we assume toxin and antitoxin production, growth-mediated dilution, and binding-unbinding kinetics of the molecules. We used a phenomenological exponential function layer that translates between concentrations of toxin and resultant growth rate, with a single parameter, α, that determines the level of toxicity.

The key similarity between our experimental and computational approaches is the existence of a threshold in the molecular network that determines the growth rate of the cell. There are many potential mechanisms for such a threshold to arise, as discussed in the Introduction. We do not claim that the mechanism implemented in the computational model is the same as the experimental model. Rather, there is an underlying fundamental interplay between growth regulation and lineage structure that we will show is conserved.

To determine the effect of the growth threshold on microcolony dynamics, we scanned the rate of toxin production, keeping antitoxin production constant. (In most natural toxin-antitoxin systems, the antitoxin is unstable. We simulated this case as well, below). The simulations were seeded with a single cell growing with excess antitoxin and permitted to grow for 100 simulation minutes before changing the toxin production rate to a positive value. After several generations of growth, we found three qualitative regimes across different toxin production rates: symmetrical growth with no or little growth arrest (toxin production rate 0-2.5 /min), a critical regime with clusters of growing and growth arrested cells (toxin production rate 3-4.5 /min), and a regime of nearly instantaneous growth arrest (toxin production rate >4.5 /min) with the colony trapped in its near-initial state. Figure 3 shows representative cases with growth rate (Fig. 3a) or toxin concentration (Fig. 3b) depicted with coloring of each cell.

**Figure 3.**
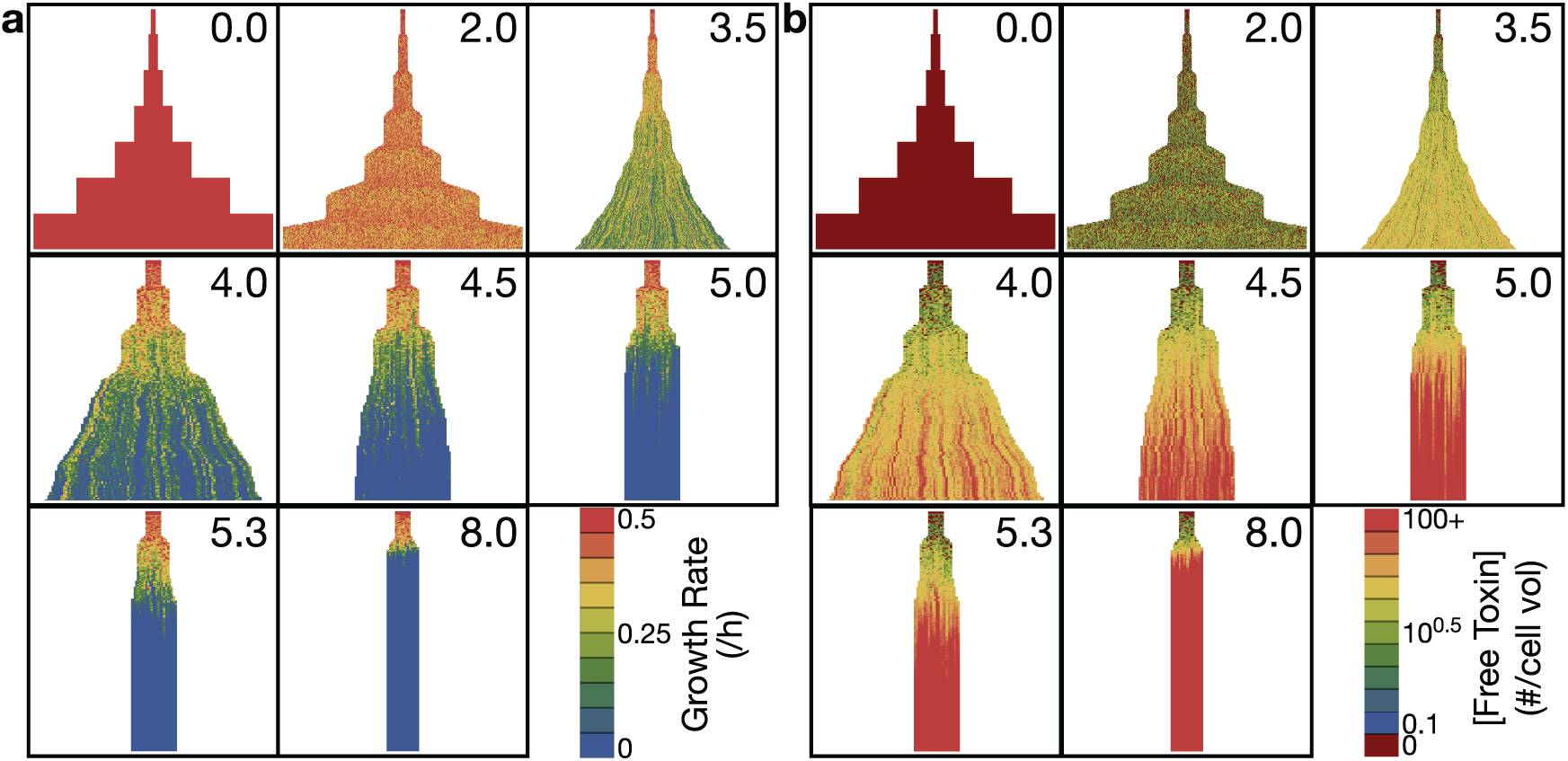
Simulated lineages over a range of toxin production rates. Time proceeds downward in each lineage and begins at the onset of toxin production (*t* = 100 h). **a.** Lineage growth rate superimposed on the lineages. **b.** Free toxin concentration superimposed on the lineage. Lineages for production rates 3.5 /min and higher are plotted with wider trajectories for visibility.

Sub-lineages of fast-growing and slow-growing cells are evident in the critical regime (with toxin production rate 5-6 /min; Fig. 3a). Lineage effects are also evident from toxin levels, where there are sublineages escaping from entry into high toxin concentrations (blue clusters in Fig. 3b). The precise time of entry into growth arrest can have a large effect on toxin levels, suggesting that growth rate is a more precise phenotype to follow for the study of lineage effects in this system.

### Lineage Dependence is Strongest in the Critical Regime

To quantitatively characterize the properties of growth transitions in our simple computational framework, we considered the fate of simulated microcolonies at 250 minutes of growth, which is shortly before the fastest growing cases begin to become computationally intractable, but after the population size is beyond the minimal requirement to be considered a microcolony. Mean population growth rates and toxin concentrations across multiple (N = 100) replicates reveal a growth-regulatable region flanked by regions of almost full growth and almost complete growth arrest (Fig. 4a). In the region where population growth is low but positive, toxin concentrations increase monotonically but non-linearly with increases in toxin production (Fig. 4a).

**Figure 4.**
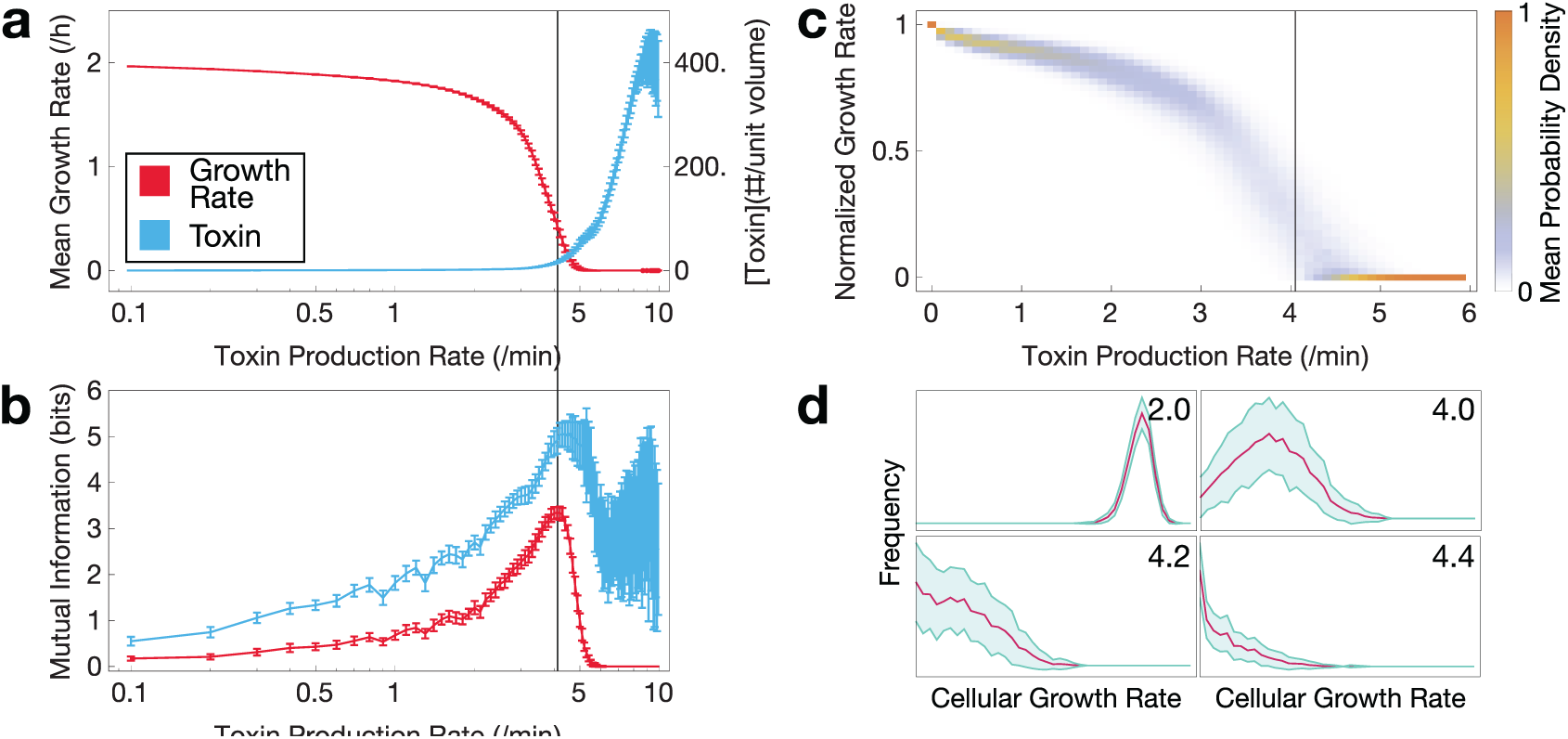
Growth, lineage information, and diversity of simulated cellular lineages at various rates of toxin production at 4 h. **a**. Average cellular growth rates (red) and toxin concentrations (blue) 150 minutes after onset of stress are proportional to toxin production rate, with distinct growth regulation regimes. Error bars indicate standard deviation. **b**. Mutual information between cell pair growth rate differences, in red (or toxin concentration difference, in blue) and their lineage distance reveals a lineage-dependent effect on cellular phenotypes near the regulatable region. **c**. Dispersion of average growth rate for low toxin production rates. Vertical bar represents the peak mutual information depicted in panel **b**. **d**. Growth rate distributions in the population at various toxin production rates as indicated. Red represents the mean frequency at a given growth rate; blue, standard deviation in the frequency.

To quantify the amount of lineage information shared by pairs of cells in their phenotypes, we calculated mutual information between phenotypic differences between pairs of cells and pairwise lineage distance. From each simulation, we sampled one pair of cells randomly to ensure independent, identically distributed samples and performed a resampling procedure 100 times to increase the confidence in our estimate. This was done for absolute growth rate differences and absolute toxin concentration differences (Fig. 4b). Various studies of have found mutual information between different points on a lattice to be indicative of a phase transition [24, 25]. While our model may not exhibit a true phase transition, our mutual information estimator reveals a similar peak for both growth rate and toxin concentrations in the critical regime, where the population growth rate is most sensitive to changes in toxin production rates.

Distributions of growth rates reveal the underlying population structure not evident from mean growth rates shown in Figure 4a. Distributions that emerge from the model include uniformly fast (Fig. 4c, top left in Fig. 4d) or slow growing (Fig. 4c, bottom right in Fig. 4d), bimodal between fast and slow growing (top right in Fig. 4d), and long-tailed with a peak at either fast (at toxin production rate 3 /min, not shown) or slow growing (bottom left in Fig. 4d).

### Fluctuating Cell Growth Dynamics in the Critical Regime

To examine a further indicator of criticality in this system, we calculated the dynamics of growing cell numbers below (toxin production rate 0-2.5 /min), near (toxin production rate 3-4.5 /min), and above the regulatable region (toxin production rate >4.5 /min) of growth rate. With toxin production well below the regulatable region, the predicted cell growth becomes equivalent to an uncoupled case where toxin has no effect on growth.

Growing cell numbers show variability between simulation replicates near the critical region (Fig. 5a). Over time, the dynamics of the mean number of growing cells approaches exponential growth at low toxin production rates, critical growth at intermediate toxin production rates (as shown in Fig. 5a), and extinction (elimination of all growth) at high toxin production rates. Mean cell numbers in critical growth show persistent oscillations that dampen as the simulated growth rates become decorrelated by noise (Fig. 5a). As toxin production approaches the critical regime, some cells accumulate high toxin and, depending on individual cellular toxin accumulation, subsets of the population will enter the exponential or extinction phase. Thus, the time required to conform to the exponential or extinction regimes is high in the critical regime, reminiscent longer relaxation times observed near critical points in other models [e.g. 26]. Autocorrelations of growing cell numbers at lag times after the onset of toxin production reveal this effect. For example, high autocorrelation around lag time 100 min in critical regime (vertical dotted line) signifies growth remaining correlated for a longer time compared to the autocorrelation at toxin production rate 3.0 /min. The presence of more than two zeroes in the absolute autocorrelations indicates the oscillatory regime (Fig. 5b).

**Figure 5.**
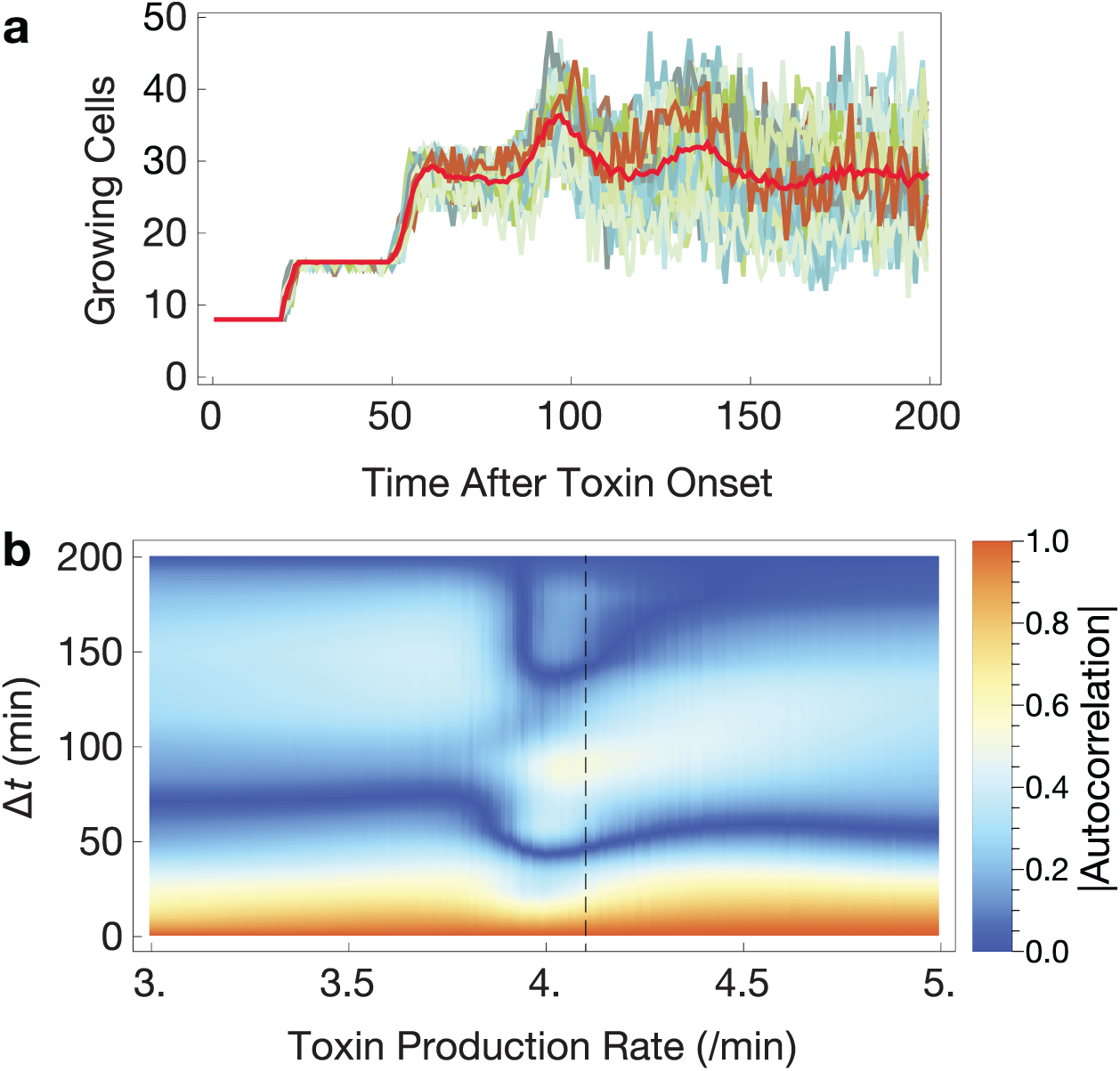
Critical slowing down of growing cell dynamics. **a**. Growing cell numbers over time in individual simulations (blue-green lines) and averaged between them (red line) reveals persistent dampening oscillations in the critical regime. **b**. Mean absolute autocorrelations near the critical regime. Δ*t*, lag time after onset of toxin production. Toxin production rates with three zeroes indicate oscillatory solutions that converge slowly to the regimes of exponential growth or extinction. Vertical dashed line indicates peak lineage-growth rate mutual information; see Fig. 4. *N* = 100 simulations for each toxin production rate.

### Attainable Levels of Phenotypic Heterogeneity Under Lineage Constraints

If lineage is capable of constraining the attainable phenotypes of offspring cells, it stands to reason that the amount of phenotypic heterogeneity attainable in a microcolony is lowered by lineage dependence in systems that generate heterogeneity by diversifying growth rates. It is difficult to generalize what constitutes meaningful diversity in growth rates; small changes may or may not be important to fitness in the long run, but the importance of the distinction between growth arrested and fast-growing cells is clear. Therefore, we used two possible definitions of meaningful diversity: in one, arbitrarily small changes in growth rate or toxin concentration are meaningful. In the other extreme, we assumed that only growing versus non-growing cells (or high versus low toxin) is a meaningful distinction.

We quantified the phenotypic heterogeneity as information entropy (base 2), binning the simulated cells according to the two definitions of diversity (Fig. 6). We calculated the maximum entropy in the fine-grained binning case by assuming each cell had a unique value. Note that the maximum entropy is extensive, decreasing with lower total cell count (Fig. 6a). In the binary case, the maximum entropy is simply 1 bit. Regardless of the definition used, the peak entropy of the population can get surprisingly close to the maximum entropy. Note that peak entropy of growth rate nearly coincides with peak mutual information between growth rate differences and lineage distance (Fig. 6, vertical line). However, entropy away from this peak sharply decreases from the maximum. In the critical regime, population heterogeneity is affected by two key factors: sensitivity of growth rate to toxin and lineage dependence. Given that we observed higher lineage dependence in the critical regime, the key question here is whether this dependence reduces the possible attainable heterogeneity in bet-hedging. The entropy plot (Fig. 6) shows that sensitivity of growth rate to toxin dominates and thus phenotypic heterogeneity is maximal at when the lineage is most structured.

**Figure 6.**
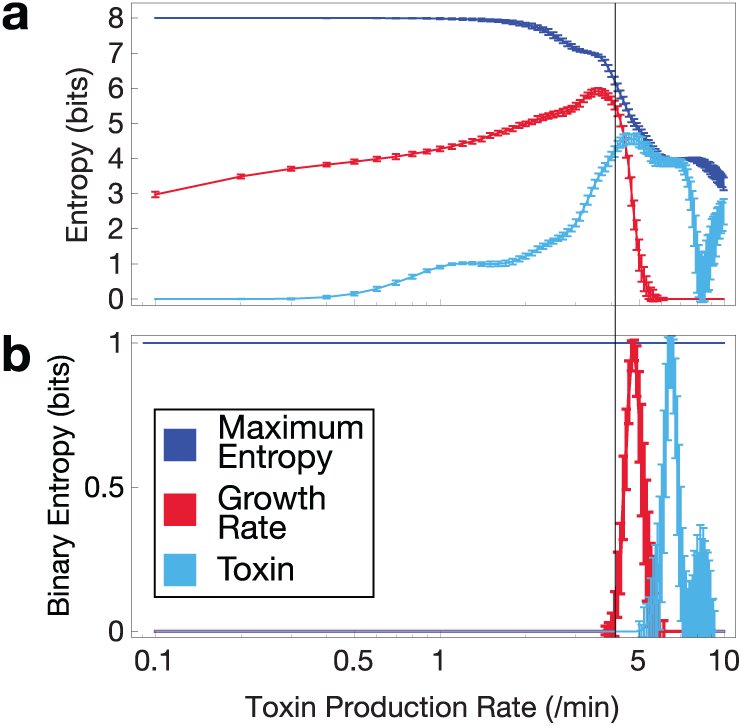
Entropy of growth rates and toxin concentrations at 250 h. Vertical line indicates the point of highest lineage-dependent mutual information between growth rate and lineage distance. **a**. Fine-grained binning. **b**. Binary binning into growing-non growing or high-low toxin concentration. Error bars indicate standard deviation.

### Growth Regulation as a Criterion for Lineage Dependence

To explore the generality of our results, we created models with variations on the original, and tested for lineage dependence.

The first set of variations test two simplifications in the primary model: stability of the antitoxin, and bursty production of the molecular species. While we regard the model to be a general threshold-based growth control mechanism, it is worthwhile to determine if a toxin-antitoxin module with unstable antitoxin qualitatively reproduces our main results. Varying the stability of the antitoxin, we indeed found the same qualitative results (Fig. S4a). Similarly, simulating bursts of gene expression producing toxin and antitoxin produced the same qualitative results (Fig. S4b).

Our next model variation was to vary the effect of growth regulation, increasing it (α=0.3 in *g*(*T*,*t*); see Methods below) and abolishing it completely (α=0 in *g*(*T*,*t*)). As expected, a larger quantitative effect of toxin preserved the main results, but shifted the toxin concentration necessary to see the lineage dependence (Fig. S4c). Abolishing growth regulation eliminated the peak in mutual information, and thus lineage dependence (Fig. S4d).

### Distributions of Growth Arrested Cluster Sizes

Large clusters of growth arrested cells could have effects on the spatial development of bacterial colonies, as daughter cells tend to be correlated in space as well. We therefore asked what growth arrested cluster size distributions arise in the region where there is high mutual information between growth rate and lineage distance. We performed 10,000 simulations each and clustered the end-point populations according to lineage neighbors having similar growth rate (with a cutoff of 0.01 /h to be considered growth arrested). Resulting clusters were pooled across simulations of the same parameter set. We present distributions of raw absolute cluster size, not normalized.

Below the critical regime, the absolute cluster size distribution is nearly exponential (Fig. 7, red line with exponential fit as gray dashed line). As the probability of growth arrest increases (with high toxin production rate), the distributions diverge from exponential to make large clusters of growth arrested cells more likely (Fig. 7). At higher toxin production rates, the distribution is bimodal between large clusters and single growth arrested cells.

**Figure 7.**
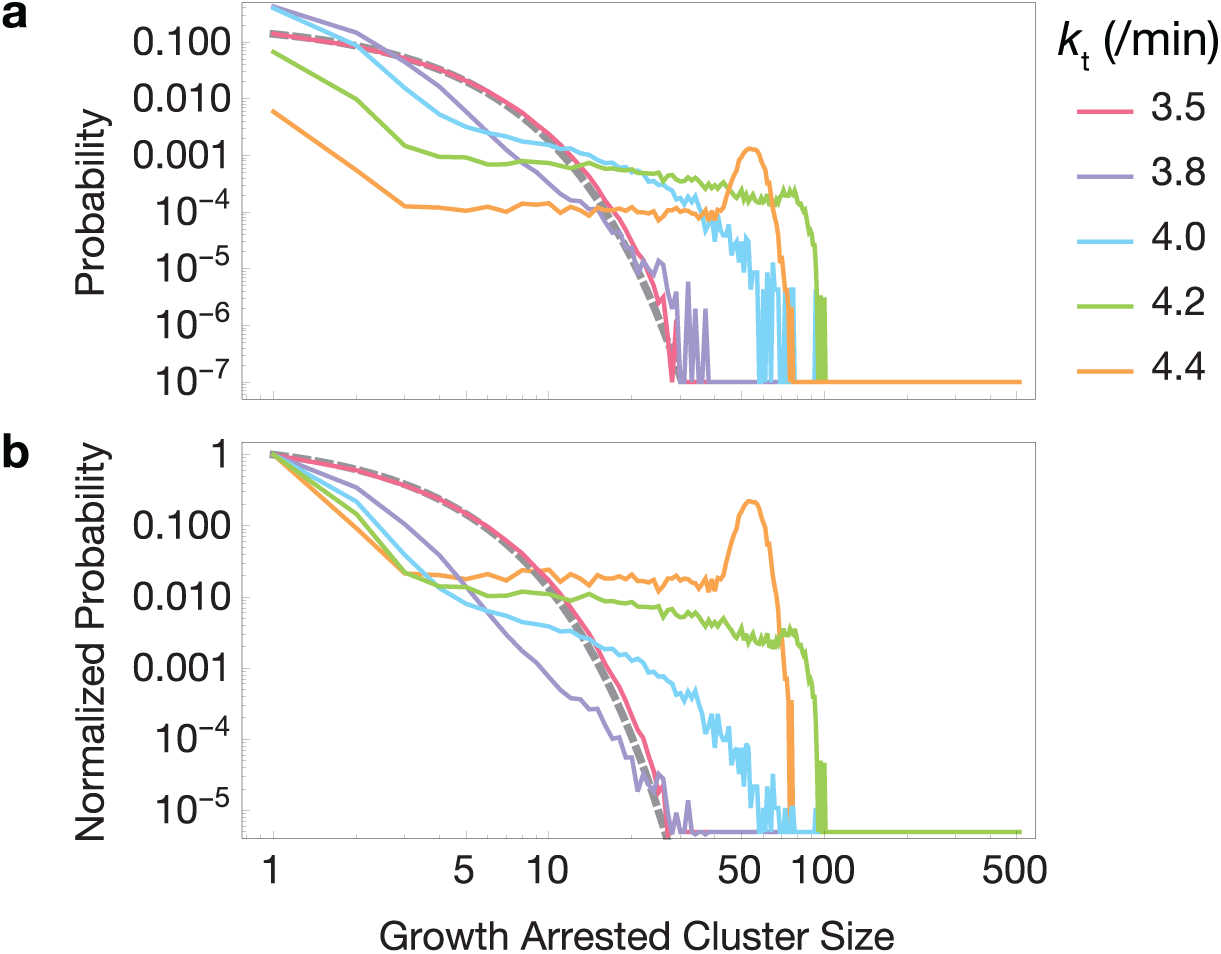
Distribution of growth arrested cluster sizes in simulated lineages. Clusters are exponentially distributed below the critical region (red line, simulation; gray dashed line, exponential fit *ae^−bc^* for cluster sizes *c*) but diverge from an exponential distribution near the critical region, eventually becoming bimodal (purple, blue, green, and orange lines). Each parameter set was simulated 10,000 times. **a**. Raw probability distributions. **b**. Probability distributions normalized to the probability of cluster size 1.

## Discussion

Regulation of growth is a central part of phenotypic control. Many factors can control growth rate, including extrinsic conditions such as starvation, and intrinsic regulators of growth that often operate with a threshold-based mechanism. Using an experimental model of threshold-based growth arrest arising from metabolic toxicity, we tracked cell growth in a bacterial microcolony with a high probability of undergoing the growth arrest transition, and a colony grown in a condition that does not display the threshold-based growth arrest. We found several large, discrete shifts in growth rate to occur at a faster timescale than our 5-minute recording intervals (Fig. 2). Quantifying the lineage dependence of cellular growth phenotype, we found that growth arrested or dead cells tend to be clustered in the lineage, as do fast-growing cells. The difference in lineage shapes between the growth arrest-prone and and non-growth arrest prone conditions is striking (Fig. 2d,h).

We therefore sought the simplest possible model of microcolony growth dynamics that reproduces the effect. Our basic model captures single-cell biochemical kinetics on one scale (microscopic) interfacing population growth dynamics on another scale (macroscopic). We found striking phenotypic lineage dependence to emerge with the following criteria: (*i*) growth rate dependence on a toxin; (*ii*) stochastic dynamics around a cellular threshold embedded within the network; (*iii*) kinetic parameters calibrated so that the population average growth rate is near the regulatable region.

As the probability of cellular transition to growth arrest increases, the mutual information between growth rate and lineage distance increases to a peak, then decreases as the simulated microcolony reaches the condition of immediate growth arrest. This transition bears a resemblance to a phase transition, with correlation of microscopic length scales peaking at the critical boundary. Here, the correlation length is in lineage space: we have assumed no traditional spatial information about the cells in the simulation.

Lineage space is a binary tree growing with extinction probability based on microscopic dynamics. Distances are modified by dynamical growth rates, which explains why a higher probability of heterogeneous growth results in structured trees. Thus, relating persister and other threshold-based growth arrest mechanisms to the established mathematics of branching processes [27, 28] is an important direction for microbial physiology.

After 100 simulated minutes we imposed a continuous rate of increased toxin production (or antitoxin degradation, in one derived model) on the developing microcolony. The constant input of more toxin created an irreversible threshold. Once a cell crosses the growth arrest threshold, there is an irreversible stoppage of growth that arises from toxin growth feedback. The growth arrest condition can then be considered an absorbing state. Continuous transitions from active to absorbing states are generically characterized by the scaling properties of critical directed percolation [29-31]. Our model qualitatively reproduces characteristics of directed percolation, including longer relaxation times near the critical region (Fig. 5) and different regimes of growth arrested cluster size distribution (Fig. 7). However, the dimensionality of the space is unclear, and may be shaped by the probability of growth arrest. Thus, we are doubtful that bet hedging quantitatively conforms to the classic criteria for directed percolation.

If lineages impart spatial structure onto growth phenotypes, then do they impose an upper limit to the level of phenotypic heterogeneity that can be attained by a microcolony? The population is most sensitive to fluctuations directly in the region with the highest lineage dependence, the latter of which appears to imply a dampening of phenotypic heterogeneity. However, multiple methods of measuring total population entropy suggest that the population can still approach the maximum total entropy in cases where growth rates are both finely-binned and binned into only two phenotypes – growing and growth arrested (Fig. 6). Heterogeneity is reduced as the population reaches either extreme of high or low average toxin level. Thus, counterintuitively, a more highly structured lineage yields a higher level of heterogeneity. Lineage plays an interesting role in determining the phenotypes of extant growing cells, but it does not appear to restrict what phenotypes can be attained.

The purely intracellular phenomena considered here allow lineage to be the only type of space considered. However, closely related cells in many conditions, such as surface-attached conditions or channels, will be physically closer together as well. In many bacterial colonies with a substantial chance of endogenous and exogenous conditions interacting to determine the growth arrest transition (such as quorum sensing), an information metric that includes components of both real space and lineage space will need to be considered.

## Methods

### Cell Culture Conditions

*E. coli* B REL606 *lacI*^−^ P_lacO1_-GFP was grown from -80° C cryogenic culture for 18 h in LB medium in a shaking incubator (37° C), acclimatized by incubating in Davis minimal medium containing either 50 mg/ml lactose (DMlac50) or 2 mg/ml glucose (DMglc2) for 24 h, and resuspended either in fresh DMlac50 or DMglc2 culture, respectively, for 3 hours before beginning time-lapse microscopy.

### Microscopy and Image Analysis

We used an Olympus IX81 inverted fluorescence microscope with an incubated imaging chamber (Olympus, Tokyo, Japan). The chamber with objective was pre-heated, bacterial cultures were added to a pre-heated CellAsic ONIX microfluidic plate (Millipore, Billerica, Massachusetts) at an approximate OD450 of 0.005, and a continuous media flow of 1 psi DMlac50 or DMglc2 was maintained for the duration of the experiment. Images in brightfield and green fluorescence (488 nm stimulation / 509 nm emission) channels were captured every 5 minutes with a 4k CMOS camera, followed by ZDC autofocus. For the DMlac50 experiment, we used a 100x oil immersion objective. Due to technical issues with the objective, we used a 60x air objective for the DMglc2 experiment. Thus, the pixel lengths of the cells between the two experiments should not be directly compared.

Images were cropped after identifying a stable microcolony originating from a single cell. We developed a semi-supervised cell tracking algorithm in Mathematica (Wolfram Research, Champaign, Illinois) with manually input cell division times and cell lengths. From this information, we reconstructed the lineage and approximated growth rates with exponential growth models. When mapping the growth rates to the lineages in Fig. 2, we approximated growth rates of cells with non-significant exponential fits using piecewise linear regression as reviewed in [32].

### Multiscale Growth Simulation Framework

To capture the minimal mechanisms necessary that recapitulate non-genetic inheritance and effects of cellular lineage, we created a multiscale growth simulation framework with individual cell agents, each containing a molecular network of interacting proteins, referred to as toxin and antitoxin, with toxin affecting cellular growth rate.

We track the simulated number of toxin and antitoxin molecules as well as cell volumes for each cell agent across time. In the next time step, *t+δt*, the number of toxin and antitoxin molecules are determined by stochastic simulation (below) and are updated for that cell. Cellular growth rates are set by a deterministic function of the toxin concentration (#/vol). The change in the volume (δ*v*) in δ*t* is determined by the amount of toxin present at that time. When cell volume doubles, the number of each molecule is distributed binomially into the two daughter cells. From that time on, the two daughter cells are labeled as different cells and are iterated in the same way. We initiate each simulation as a single cell with no toxin and allow growth for a few generations (100 minutes) before applying toxin production rate (or antitoxin degradation rate) of a given quantity. The primary purpose of this model is to capture the qualitative effect of the growth arrest threshold, so several important details about the biophysics of kinetics in growing cells were omitted, such as the effects of chromosome replication and the volume dependence of bimolecular stochastic reaction propensities.

### Estimation of Mutual Information from Simulated Lineages

We sought to develop a sampling methodology to ensure independent, identically distributed samples from lineage simulations to estimate the mutual information between lineage distance *d* and phenotypic differences between pairs of cells *φ*. Phenotypic differences (*φ*) could be growth rate or intracellular toxin concentration. To do so, we performed 100 independent simulations in each condition, and randomly drew a single pair of cells from each lineage. Our estimate of mutual information was calculated from the resulting distribution of i.i.d. samples: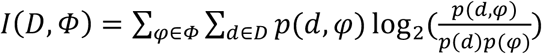. A more accurate estimate of absolute mutual information may extrapolate to an infinite sample size. In our case, the relative mutual information between different locations in parameter space suffices to demonstrate the existence of a strong lineage dependence for certain parameter ranges. To estimate the uncertainty of our relative mutual information estimate, we resampled 100 cell pairs with replacement and present the resulting mean ± standard deviation. Entropy was calculated by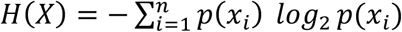, where *P*(*x*_*i*_)repre-sents the probability mass function of a discrete variable *X*. *X* could be growth rate or toxin concentration.

### Stochastic Toxin-Antitoxin Threshold Model

We considered a simple network consisting of three variables: toxin, antitoxin and toxin-antitoxin bound complex. Possible reaction events are synthesis of toxin and antitoxin, and binding and unbinding between toxin and antitoxin molecules. The reaction scheme for the basic model is:

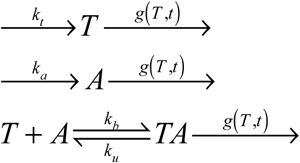

The parameter *k*_*t*_ is the toxin production rate varied in the simulations. Antitoxin production parameter, *k*_*a*_, is kept constant (*k*_*a*_ = 4.2 /min) to allow the production ratio of toxin and antitoxin to be changed. Growth-mediated loss is implemented through *g*(*T*, *t*) which is a function of the cell volume in the algorithm (below). Parameters *k*_*b*_ and *k*_*u*_ are binding and unbinding rates; *k*_*b*_ = 0.1 and *k*_*u*_ = 0.1 throughout. In the most basic model, each species is considered long-lived on the timescales of the simulation, so we do not consider any additional degradation processes. Variations on this model are discussed in Results.

The relationship between toxin concentration and cellular growth rate, the most phenomenological part of the framework, captures the interface between molecular and population dynamic scales. We reasoned that, while some random factors may reduce or increase the effect of toxin, the generality with which toxin affects global protein synthesis rates [11, 33-37] means that many stochastic effects will cancel, resulting in a nearly deterministic relationship. Because toxin levels generally halt ongoing processes without significant delay [38-41], we approximated the effect of a given toxin level to be instantaneous. This assumption is supported by our experimental results, which show shifts in growth rate faster than the 5 minute intervals measured (Fig. 2). We thus constructed a deterministic function to reflect the functional dependence of growth on toxin concentrations: *g*(*T*, *t*) = *λe*^−*αT*(*t*)/*Ω*(*t*)^, where *α* is a parameter that represents the toxicity of the toxin, *T*. We used α = 0, 0.1 and 0.3 to represent cases with no toxicity, moderate toxicity, and high toxicity, respectively. Python scripts are given in S1-S3 Model.

### Simplified Computational Model of Binomial Inheritance

To illustrate the effects of growth arrest on distributions of growth-modulating cytoplasmic contents (**Fig. 1**), we created a simplified computational model with constant production, constant sub-threshold generation times, and binomially distributed molecular contents between two daughter cells. One simulation for each initial condition was run for 12 generations, with 10 molecules produced per generation, and a growth arrest threshold of 20 molecules. Initial conditions were 0, 10, 20, or 30 molecules. A second case with no threshold was simulated with the same parameters and initial conditions. The Mathematica code is given in S4 Model.

### Deterministic Molecular-Scale Model as a Basis for Growth Feedback

The exact functional dependency of growth on toxin is unknown. In our stochastic simulation framework, we considered an exponential dependence of growth on toxin. Fig. 1b depicts a deterministic model of toxin growth feedback by a free toxin as follows: 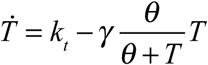, where *k*_*t*_ is the toxin production rate, *γ* is the maximum growth rate, and *θ* determines the toxicity level of the toxin. We chose the Hill form for the deterministic model because it has a closed-form steady state. The steady state is 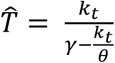 When *k*_*t*_/*θ* > *γ*, there is no steady state at this scale and the containing cell is expected to enter growth arrest. This simple model demonstrates the basis for growth feedback-induced growth arrest in a single cell. For Fig 1b, parameters are: *k*_*t*_ = 4.2 /min, *γ* = 0.023, and *θ* = 100 molecules. We note that the basic growth arrest threshold effect readily emerges in both Hill and exponential model forms, and likely a variety of other mathematical forms.

## Acknowledgments

We acknowledge Sheng Chen, Dominique Chu, Eric J. Deeds, and Uwe Täuber for useful conversations. Microscopy experiments were performed in the Microscopy and Analytical Imaging Laboratory at the University of Kansas. This project was supported by Institutional Development Awards (IDeA) from the National Institute of General Medical Sciences of the National Institutes of Health under Grant Numbers P20GM103418 and P20GM103638. The content is solely the responsibility of the authors and does not necessarily represent the official views of the National Institute of General Medical Sciences or the National Institutes of Health.

## Supporting Information Captions

**S1 Video. *E. coli* microcolony undergoing frequent growth arrest.** Time-lapse fluorescence microscopy of a cell lineage of *Escherichia coli* B. REL606 *lacI*^−^ P_lacO1_-GFP in DMlac50. Cells are tracked and measured as indicated. Numbers represent time (minutes) after the first frame. Experimental details are given in Methods.

**S2 Video. *E. coli* microcolony growing without the growth arrest threshold.** Time lapse fluorescence microscopy of a cell lineage of *Escherichia coli* B. REL606 *lacI*^−^ P_lacO1_GFP in DMglc2. Cells are tracked and measured as indicated. Numbers represent time (minutes) after the first frame. Experimental details are given in Methods.

**S1 Figure**. **Growth trajectories for all cells in the microcolony depicted in Figure 2.**

**S2 Figure**. **Probability distribution of lineage distance (time since most recent common ancestor) for the experimental lineage.** All cells (**a**), only non-growth-arrested cells (**b**), and only growth-arrested cells (**c**) in the lineage shown in Figure 2**d**. *p* < 0.01 for growth-arrested cells to not to have lower lineage distances versus either of the other two groups (one-tailed Mann-Whitney U test).

**S3 Figure. Non-exponential cell length trajectories.** Lengths of cells between divisions were tested for a significant fit to an exponential growth model in the growth arrest-prone condition. These cases failed the significance test with a Bonferroni-adjusted *α* = 0.05 (adjusted value = 0.000424).

**S4 Figure. Computational model extensions preserve the central results. a**. Altering toxin degradation rates to represent the precise mechanism of toxin-antitoxin systems. **b**. Altering toxin and antitoxin production so that they are bursty with a telegraph (ON-OFF) model. **c**. Increasing toxicity with parameter *α* = 0.3. **d**. Eliminating growth feedback (*α* = 0) eliminates the peak of mutual information along with the lack of macroscopic growth regulation.

**S1 Model. Python script for simulating lineages with stochastic simulation of the intracellular toxin-antitoxin system.**

**S2 Model. Python script for simulating lineages with stochastic simulation of the intracellular toxin-antitoxin system with bursty telegraph model of toxin and antitoxin production.**

**S3 Model. Python script for simulating lineages with stochastic simulation of the intracellular toxin-antitoxin system with fast degradation of the antitoxin.**

**S4 Model. Simplified computational model of binomial inheritance Mathematica file.**

**S1 Data. Data used to generate plots in Figure 3.**

